# Maintained representations of the ipsilateral and contralateral limbs during bimanual control in primary motor cortex

**DOI:** 10.1101/2020.03.30.015842

**Authors:** Kevin P. Cross, Ethan A. Heming, Douglas J. Cook, Stephen H. Scott

## Abstract

Primary motor cortex (M1) almost exclusively controls the contralateral side of the body. However, M1 activity is also modulated during ipsilateral body movements. Previous work has shown that M1 activity related to the ipsilateral arm is independent of the M1 activity related to the contralateral arm. How do these patterns of activity interact when both arms move simultaneously? We explored this problem by training two monkeys (male, Macaca mulatta) in a postural perturbation task while recording from M1. Loads were applied to one arm at a time (unimanual) or both arms simultaneously (bimanual). We found 83% of neurons were responsive to both the unimanual and bimanual loads. We also observed a small reduction in activity magnitude during the bimanual loads for both limbs (25%). Across the unimanual and bimanual loads, neurons largely maintained their preferred load directions. However, there was a larger change in the preferred loads for the ipsilateral limb (~25%) than the contralateral limb (~9%). Lastly, we identified the contralateral and ipsilateral subspaces during the unimanual loads and found they captured a significant amount of the variance during the bimanual loads. However, the subspace captured more of the bimanual variance related to the contralateral limb (97%) than the ipsilateral limb (66%). Our results highlight that even during bimanual motor actions, M1 largely retains its representations of the contralateral and ipsilateral limbs.

**Significance Statement:** Previous work has shown that primary motor cortex (M1) reflects information related to the contralateral limb, its downstream target, but also reflects information related to the ipsilateral limb. Can M1 still reflect both sources of information when performing simultaneous movements of the limbs? Here we use a postural perturbation task to show that M1 activity maintains a similar representation for the contralateral limb during bimanual motor actions, while there is only a modest change in the representation of the ipsilateral limb. Our results indicate that two orthogonal representations can be maintained and expressed simultaneously in M1.

## Introduction

Motor cortex is primarily involved with controlling the contralateral side of the body. Output projections from motor cortex principally target muscles for the contralateral limb (Cheney and Fetz, 1980; Dum and Strick, 1996; Brosamle and Schwab, 1997; Lacroix et al., 2004; Rosenzweig et al., 2009; Kuypers, 2011; Soteropoulos et al., 2011) and micro-stimulation in motor cortex elicits mainly contralateral limb movements (Montgomery et al., 2013). However, activity in motor cortex is modulated by movements with either the ipsilateral or contralateral limbs (Donchin et al., 1998; Kermadi et al., 1998; Cramer et al., 1999; Ganguly et al., 2009; Diedrichsen et al., 2013; Berlot et al., 2019). Neural recordings indicate ~50% of neurons that are active for contralateral limb movements are also active for ipsilateral limb movements (Steinberg et al., 2002; Cisek et al., 2003; Heming et al., 2019). Ipsilateral-related activity also exhibits broad tuning to reach direction (Steinberg et al., 2002; Cisek et al., 2003) and applied loads (Heming et al., 2019).

A largely unexplored question is how motor cortex represents the limbs during bimanual movements. Many neurophysiological investigations of bimanual movements have focused on premotor areas, such as dorsal premotor and supplementary motor cortex (Tanji et al., 1987, 1988; Donchin et al., 1998; Kermadi et al., 2000; Willett et al., 2020). During unimanual reaches, these areas exhibits similar tuning for the contralateral and ipsilateral limbs (Steinberg et al., 2002; Cisek et al., 2003) with overlapping subspaces (~50%) for the contralateral- and ipsilateral-related activity (Willett et al., 2020). During bimanual motor actions, the contralateral-related activity is largely unchanged, whereas the ipsilateral activity is reduced by ~50% (Rokni et al., 2003; Willett et al., 2020). It has been hypothesized that the suppression of the ipsilateral representation and its decoupling from the contralateral representation reduces its interference on the descending contralateral motor commands during bimanual control (Rokni et al., 2003; Willett et al., 2020).

However, it is unclear if a similar change and suppression of the ipsilateral-related activity would occur in primary motor cortex (M1). During unimanual movements, M1 has decoupled representations for the contralateral and ipsilateral limbs as neurons are tuned independently for each arm (Cisek et al., 2003; Heming et al., 2019) and contralateral- and ipsilateral-related activities occupy orthogonal subspaces (Ames and Churchland, 2019; Downey et al., 2019; Heming et al., 2019). Thus, M1 could maintain its representations of each limb across unimanual and bimanual movements as the representations are already decoupled.

We explored this hypothesis by training monkeys in a postural perturbation task. Monkeys performed this tasking using only one arm at a time (unimanual) and using both arms simultaneously (bimanual). We found almost all neurons active during unimanual loads were also active for bimanual loads, and vice versa. There was a small reduction in the magnitude of activity related to both arms during the bimanual loads. We also found neurons largely maintained their preferred load direction across the unimanual and bimanual loads, with a stronger relationship for the contralateral-related activity than the ipsilateral-related activity. Lastly, the contralateral and ipsilateral subspaces identified during the unimanual loads captured a significant amount of variance for the bimanual loads.

## Methods

### Animals and apparatus

Two male non-human primates (Macaca mulatta, weight ~15kg) were trained to place their arms into an exoskeleton robot (Kinarm, Kingston, Canada (Scott, 1999)) and perform a postural perturbation task similar to our previous work (Herter et al., 2009; Pruszynski et al., 2014; Heming et al., 2019). At the start of each trial, a target appeared (0.8cm diameter, red for right, blue for left, luminance matched) that was placed in front of the shoulder with a starting joint position of 30° at the shoulder and 90° at the elbow. The monkey held their hand inside the target for 500-1000ms, after which a load was applied by the exoskeleton that displaced the hand from the target. The monkey had 1000ms to return their hand to the target and hold within the target for 1000-1500ms to receive water reward. On a given trial, the monkey performed this task with only one hand (Figure 1A,B unimanual contexts contra-only, ipsi-only Figure 1A,B) or both hands at the same time (Figure 1C,D bimanual contexts mirror, opposite). The appearance of one or two targets at the start of the trial cued the monkey about whether one hand or both hands were required. Within a block, all unimanual and bimanual trials were randomly interleaved, and monkeys completed a minimum of 10 blocks.

**Figure 1:**
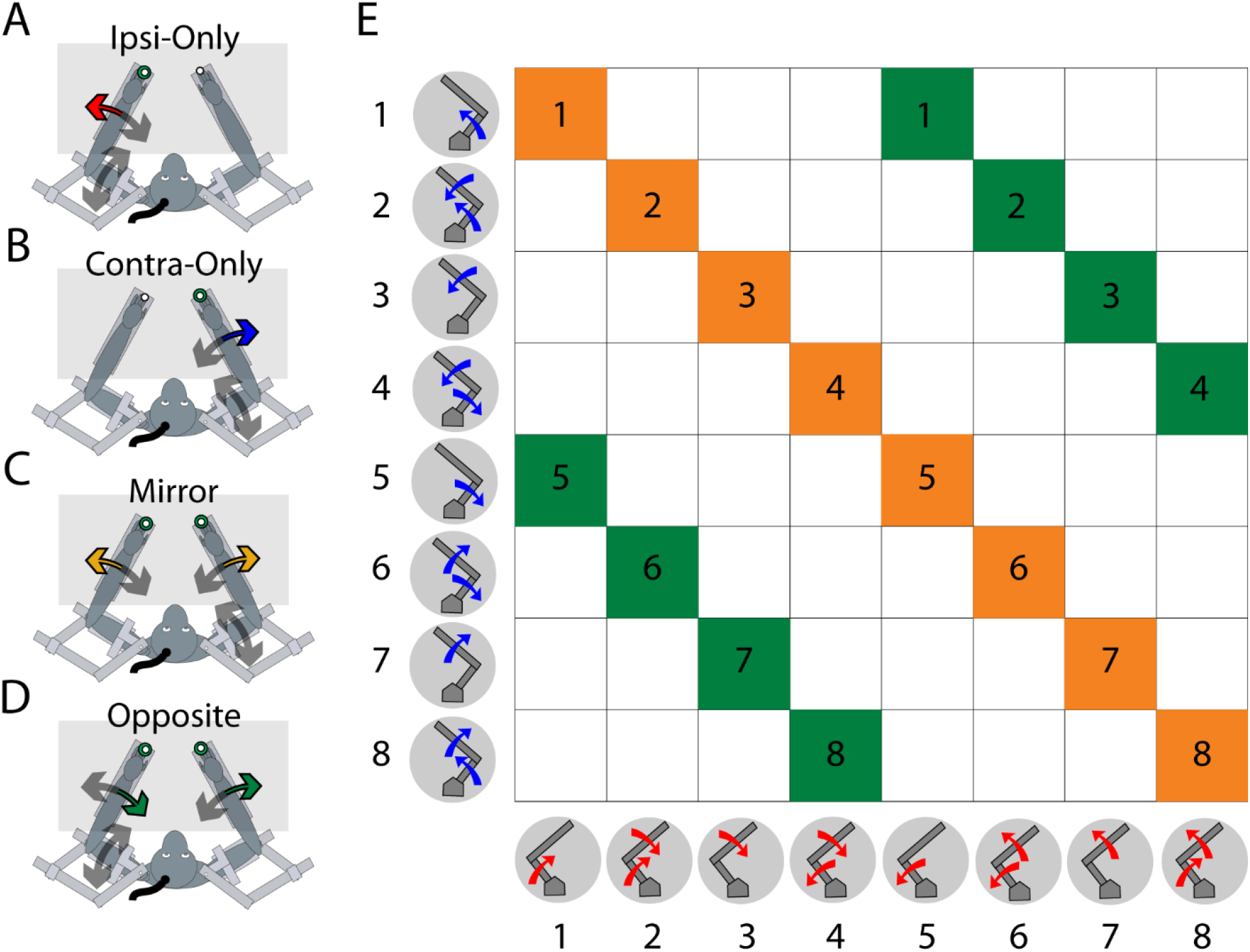
Experimental set-up. Monkeys were trained in a postural perturbation task using their left and right limbs. We included trials where loads were applied to only one limb at a time (A-B) or to both limbs simultaneously (C-D). E) A table showing all possible load combinations. We selected only combinations where the contralateral and ipsilateral loads were equal (yellow squares, mirror) or where the loads were equal in magnitude but opposite in sign (green squares, opposite).

Loads consisted of flexion and extension torques applied to the shoulder and/or elbow joints. Eight combinations were used, including four single-joint torques (elbow extension (EE), elbow flexion (EF), shoulder extension (SE), and shoulder flexion (SF)) and four multi-joint torques (SF/EF, SF/EE, SE/EF, SE/EE). For Monkey P, single-joint torques consisted of ±0.20Nm torques (+=flexion load, -=extension load), whereas multi-joint torques consisted of ±0.14Nm torques applied to both joints. Monkey M completed this task with two different torque magnitudes, a large and a small load set. The large/small load set included single-joint torques of ±0.30/0.20Nm and multi-joint torques that consisted of ±0.24/0.14Nm torques applied to both joints.

For the bimanual loads, it was not feasible to test all possible torque combinations between the two arms (Figure 1E). Instead, we focused on load combinations that were mirror symmetric across both arms (orange squares, e.g. contralateral SF/EE, ipsilateral SF/EE) and load combinations that were equal magnitude but opposite direction (green squares, e.g. contralateral SF/EE, ipsilateral SE/EF).

### Neural and kinematic recordings

Monkeys had Utah Arrays (96-channel, Blackrock Microsystems, Salt Lake City, UT) implanted into the arm region of M1. Neural signals were digitized by a 128-Channel Neural Signal Processor (Blackrock Microsystems, Salt Lake City, UT) at 30kHz. An offline spike sorter (Plexon) was used to manually isolate units and we only used well-isolated single units.

For Monkey P, neural signals were recorded in three sessions spaced approximately 4 months apart. For Monkey M, when performing the task with the small loads, neural signals were recorded from two sessions spaced 3 months apart. When performing the task with the large loads, Monkey M was unable to complete all 10 blocks in one recording day. Instead, Monkey M completed the 10 blocks over the course of 2 or 3 consecutive days, yielding one session. We only included single units we could isolate consistently across the recording days and had qualitatively similar spike waveforms and inter-spike interval histograms. Two sessions were collected that were spaced 4 months apart.

Neurons across all recorded sessions for a given monkey were treated as independent and pooled. Previously, we have estimated that <5% of neurons would have overlapped between sessions that are spaced out by 3 months (Heming et al., 2019). Furthermore, Fraser and Schwartz, (2012) found only a few neurons could be tracked for >3 months on an array.

Joint angles, velocities, and accelerations were also recorded by the Neural Signal Processor at 1kHz.

### Data and statistical analysis

#### Kinematic analysis

Kinematic signals were low-pass filtered at 10Hz using a 3^rd^-order Butterworth filter. We quantified the integrated and maximal hand speed over the first 300ms after the perturbation (perturbation epoch), as well as the exact hand speed at the 300ms time point. Statistical significance was assessed using a one-way ANOVA with load context as a factor (levels: contra-only, ipsi-only, mirror, opposite). Post hoc Tukey-Kramer tests were used to assess significance between levels.

#### Spike train and time epochs

The instantaneous activity of a neuron was estimated by convolving the spike time stamps with a kernel approximating a post-synaptic potential (1ms rise and 20ms fall, Thompson et al., 1996). Activity in the perturbation epoch was calculated by aligning to the load onset and averaging across trials for the first 300ms. Steady-state activity was calculated by aligning to the load offset at the end of the trial and averaging across trials for the 1000ms that preceded the load offset.

#### Dynamic range

During the perturbation epoch, we calculated the mean activity during the epoch for each load combination, creating eight separate values for each context. The difference between the largest and smallest mean activity within a context was defined as the dynamic range. An identical procedure was used to calculate the dynamic range in the steady-state epoch. A paired t-test was used to compare the activities across neurons between the bimanual loads and the appropriate additive models.

#### Linear model fits

The mean activities for each neuron were regressed onto the applied torques to estimate tuning and magnitude. For each neuron, separate 8×1 arrays were constructed that contained the contra-only (*fr_Contra_*) and ipsi-only (*fr_Ipsi_*) activities. The mean activity of each array was subtracted and fit with the following equations

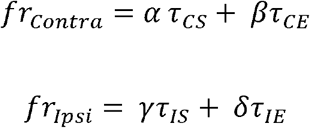

Where *τ_CS_, τ_CE_* are 8×1 arrays containing the torques applied to the contralateral shoulder and elbow joints, respectively, and *τ_IS_, τ_IE_* are 8×1 arrays containing the torques applied to the ipsilateral shoulder and elbow joints, respectively. The *α, β, γ, δ* are scalar fit parameters. For the contralateral torques, the activity magnitude of a neuron was calculated by 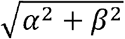, and its preferred direction was calculated as 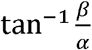. Similar formulas were used for the ipsilateral torques using the γ and *δ* fit parameters.

For the bimanual data, regressing the mirror and opposite activity on to the applied loads separately resulted in the contralateral loads being collinear with the ipsilateral loads. We mitigated this problem by concatenating the activity for the mirror (*fr_Mirror_*) and opposite (*fr_opposite_*) contexts into a 16×1 array (*fr_Bimanual_*) The mean activity of the array was subtracted and fit with the following equation

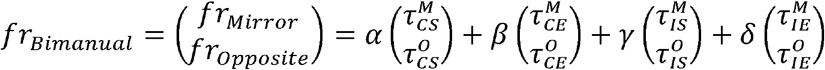

Where 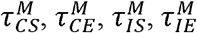 are the shoulder and elbow torques applied to the contralateral and ipsilateral limbs for the mirror loads, and 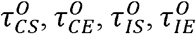 are the shoulder and elbow torques applied to the contralateral and ipsilateral limbs for the opposite loads. Note, in our experiment 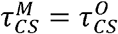 and 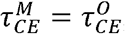 whereas 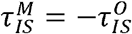 and 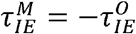.

However, by using both the mirror and opposite activities the estimated fit parameters were less affected by sampling error than the equivalent unimanual fit parameters. This was a problem for comparing activity magnitudes between contexts as higher sampling error will overestimate activity magnitude (Willett et al., 2020). Consider an example where we estimate *α* with some sampling error *η* such that 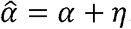. For simplicity, we assume *β* and 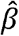 are zero, though this is not necessary. Calculating the magnitude results in 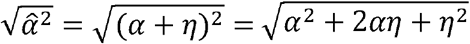. Since *η^2^*> 0 this introduces a positive bias in our estimate of the magnitude. Note, the term *2αη* can be negative, thus reducing the impact of *η^2^*. However, in simulations and our data, we still found a positive bias in the activity magnitudes.

We minimized this bias by randomly sampling half of the trials from the mirror and opposite contexts. We then trial-averaged across these samples and completed our analysis described above. We repeated this 1000x and calculated the average magnitude and preferred load direction for each neuron.

#### Change of Tuning

A neuron’s change in tuning was defined by the difference between its preferred directions for the unimanual and bimanual contexts. We constructed a distribution reflecting the change of tuning across the population of neurons. We quantified how unimodal this distribution was using the Rayleigh unimodal coefficient (R coefficient, Batschelet, 1981). We compared our results with a null distribution that randomly shuffled the neurons’ preferred directions and calculated the resulting change in angle (“Shuffle”). The R coefficient was then calculated, and the procedure was repeated 1000 times. We also generated a distribution that compared the tuning change expected from independent samples within a load context (“Within-Context”). We evenly split the contra-only trials into two separate groups. We then calculated the change in tuning between these groups by using the same procedure as above. Probability values were calculated by findings the number of R coefficients from the shuffle and within-context distributions that were greater than and less than the empirical R coefficient, respectively. We repeated this 1000 times. A similar calculation was done using the ipsi-only trials.

#### Nonlinear modeling and AIC

We also fit the bimanual activity with models that included nonlinear interaction terms between the contralateral and ipsilateral torques

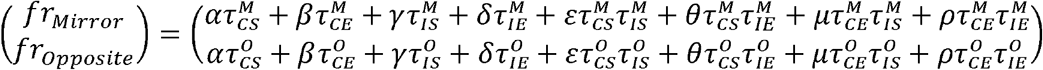

Where ε, θ, μ, *p* are scalar fit parameters. We used Akaike’s Information Criteria (AIC) to compare the linear and nonlinear models, which balances model complexity with performance (Burnham and Anderson, 2004). Given the small number of samples (16) relative to the number of parameters in each model (linear 4, interaction 8) we applied a small sample correction to the AIC.

#### Joint optimization

We identified the contralateral and ipsilateral subspaces using a joint optimization method that we have used previously (Elsayed et al., 2016; Heming et al., 2019). Briefly, this optimizer seeks a set of projections for the contralateral and ipsilateral activities that maximized the amount of variance captured while constrained to keep the projections orthogonal with respect to each other (Elsayed et al., 2016). We calculated the alignment index to quantify how well these axes aligned with the bimanual data.

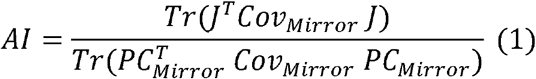

Where *Cov_Mirror_* is the mirror covariance matrix, *PC_Mirror_* is the top-ten principle components for the mirror activity. J is the top five contralateral and ipsilateral projections concatenated yielding ten projections. The alignment index can range from 0, indicating no overlap, to 1, indicating complete overlap. Simply, this metric reflects the ratio between the total amount of variance captured by J with the amount of variance captured by the top-ten mirror principle components (i.e. the most variance any 10 linear projections could capture). A similar method was used to calculate the alignment index for the opposite activity and additive models. A null distribution was generated by randomly sampling subspaces that are biased by the data covariance matrix, as previously described (Elsayed et al., 2016). Probability values were calculated by findings the number of alignment indices from the null distribution that were greater than the empirical alignment index.

## Results

### Kinematic Results

We trained monkeys to perform a postural perturbation task where loads were applied to either limb only (unimanual context), or both limbs, simultaneously (bimanual context). Monkey P was able to easily complete this task with an average success rate of 89%. Monkey M struggled with this task when the load magnitudes were 0.3Nm (“large loads”) with an average success rate of 51%. In particular, Monkey M struggled with the multi-joint bimanual loads, a problem also observed in a similar task with humans (Omrani et al., 2013). As a result, we also had Monkey M complete the same task using load magnitudes of 0.2Nm (“small loads”) in a separate set of recording sessions. With the small loads, Monkey M had an average success rate of 87%.

Figure 2A shows Monkey P’s left (ipsilateral) hand paths for all load combinations and contexts. For the first 300ms after the load onset (colored circles), the hand trajectories were similar regardless of whether the ipsilateral loads were applied without (ipsi-only) or with (mirror and opposite) an accompanying contralateral load. In contrast, when only contralateral loads (contra-only) were applied, there was little movement observed in the left hand. Similarly, Figure 2B shows the right (contralateral) hand for all load combinations and contexts. Contralateral loads evoked similar hand trajectories when accompanied without and with an ipsilateral load, whereas little motion was observed when only ipsilateral loads were applied. Examining the hand speed (Figure 2C and 2D) revealed similar observations.

**Figure 2.**
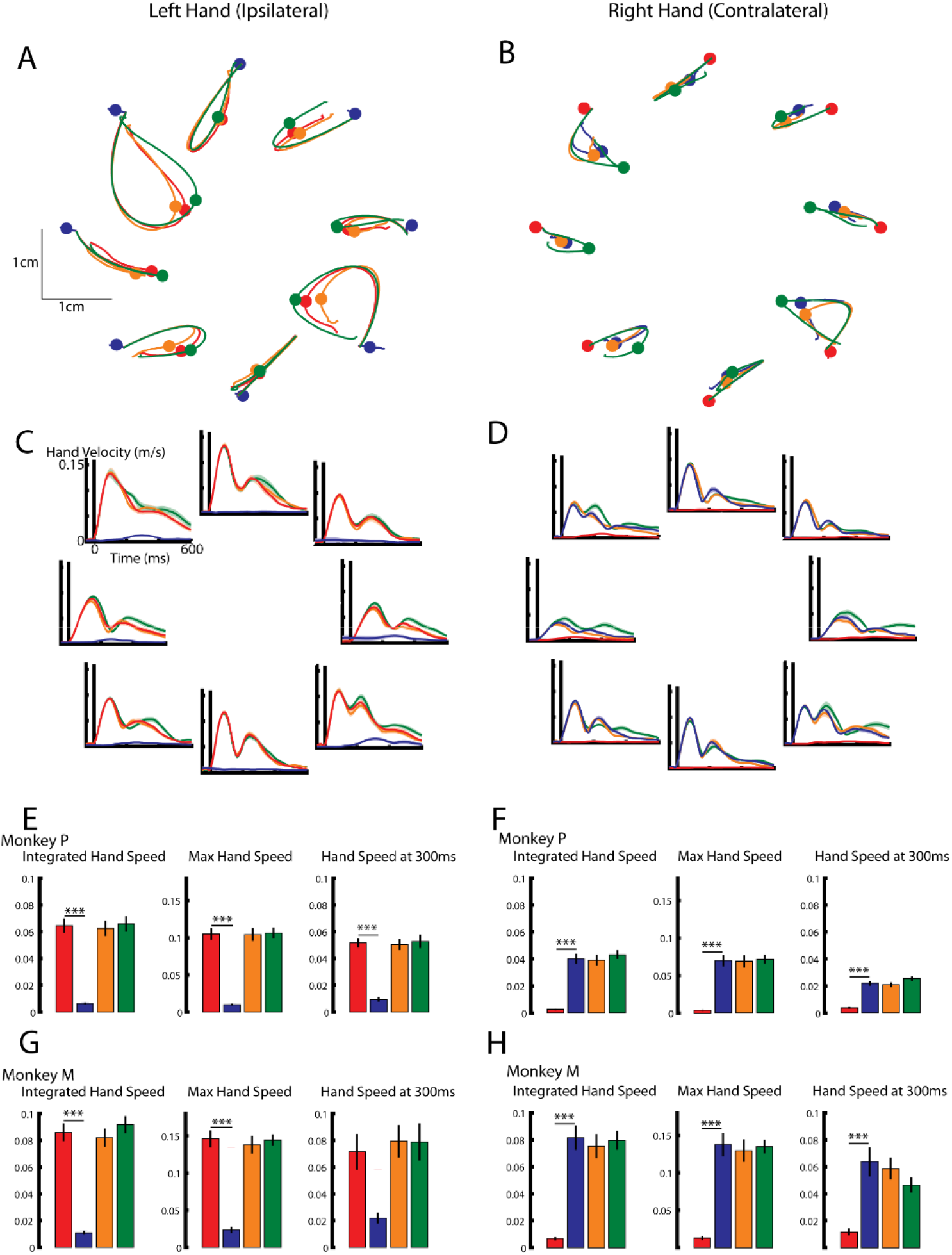
A-B, Hand paths for the left and right hand of Monkey P. Blue traces for perturbations to the contralateral limb only, red for perturbations to the ipsilateral limb only, orange and green are contralateral and ipsilateral perturbations that are mirror and opposite, respectively. Circles indicate the 300ms mark on the hand trajectory. C-D) Left and right hand speeds for each perturbation type from Monkey P. E) For the left hand of Monkey P, the mean across load combinations for the integrated hand speed, maximum hand speed, and hand speed at 300ms. A one-way ANOVA with context as a factor revealed a significant main effect for the integrated hand speed (F(3,28)=35 p<0.001), maximum hand speed (F(3,28)=48 p<0.001) and the hand speed at 300ms (F(3,28)=30 p<0.001). F) Same as E) for the right hand of Monkey P. A significant main effect was found for the integrated hand speed (F(3,28)=35 p<0.001), maximum hand speed (F(3,28)=26 p<0.001) and the hand speed at 300ms (F(3,28)=41 p<0.001). G) Same as E) except for Monkey M. A significant main effect was found for the integrated hand speed (F(3,28)=42 p<0.001), maximum hand speed (F(3,28)=41 p<0.001) and the hand speed at 300ms (F(3,28)=6 p<0.001). H) Same as F) except for Monkey M. A significant main effect was found for the integrated hand speed (F(3,28)=24 p<0.001), maximum hand speed (F(3,28)=27 p<0.001) and the hand speed at 300ms (F(3,28)=10 p<0.001). E-H) Post hoc Tukey-Kramer tests were used to compare either the unimanual ipsilateral loads (E,G) with the other three contexts or the unimanual contralateral loads with the other three contexts (F,H). *** p<0.001. All p values were Bonferroni corrected with a factor of three.

We calculated the integrated hand speed over the first 300ms for all load contexts (Figure 2E for Monkey P and Figure 2G for Monkey M large loads). For the left hand, a one-way ANOVA with load context as a factor revealed a significant main effect for both monkeys (Monkey P: F(3,28)=35 p<0.001; Monkey M: F(3,28)=42 p<0.001). Post hoc analysis confirmed that contra-only loads evoked smaller hand motion in both monkeys (left columns Figure 2E,G). Similar results were found when we examined the maximum hand speed within the first 300ms (center column), as well as the hand speed at 300ms (right column).

For the right hand, a one-way ANOVA revealed a significant main effect for the integrated hand speed for both monkeys (Monkey P: F(3,28)=35 p<0.001; Monkey M: F(3,21)=24 p<0.001). Post hoc analysis confirmed that ipsi-only loads evoked smaller hand motion in both monkeys (Figure 2F,H). Similar results were found when we examined the maximum hand speed within the first 300ms (center column), as well as the hand speed at 300ms (right column). Similar results were also found when we examined Monkey M’s kinematics for the smaller loads (data not shown).

### Neural Recordings

We recorded 92 neurons from Monkey P. From Monkey M, we recorded 66 neurons with the large loads and 78 neurons during the small loads. For Monkey M, we pooled all neurons (144) recorded for the large and small loads as our findings were similar when we analyzed each group separately.

Figure 3A shows the activity of an example neuron when ipsi-only and contra-only loads were applied (top panels). For simplicity, we only present the neuron’s activity for two of the multi-joint loads (SF/EE light colours, SE/EF dark colours). For both contexts, this neuron displayed clear selectivity for the loads, with greater activity during ipsi-only loads for SE/EF (left panel), and greater activity during contra-only loads for SF/EE (right panel). However, for the mirror context this neuron exhibited little selectivity for the loads (middle left panel). For comparison, we calculated the expected mirror activity if it simply reflected the addition of the ipsi-only and contra-only activities (additive mirror model, bottom left panel). The additive mirror model also showed little selectivity for the loads. For the opposite context, this neuron exhibited clear selectivity for the loads (middle right) and was qualitatively similar to the equivalent additive model (bottom right panel). Figure 3B and C show the activities for two additional example neurons.

**Figure 3:**
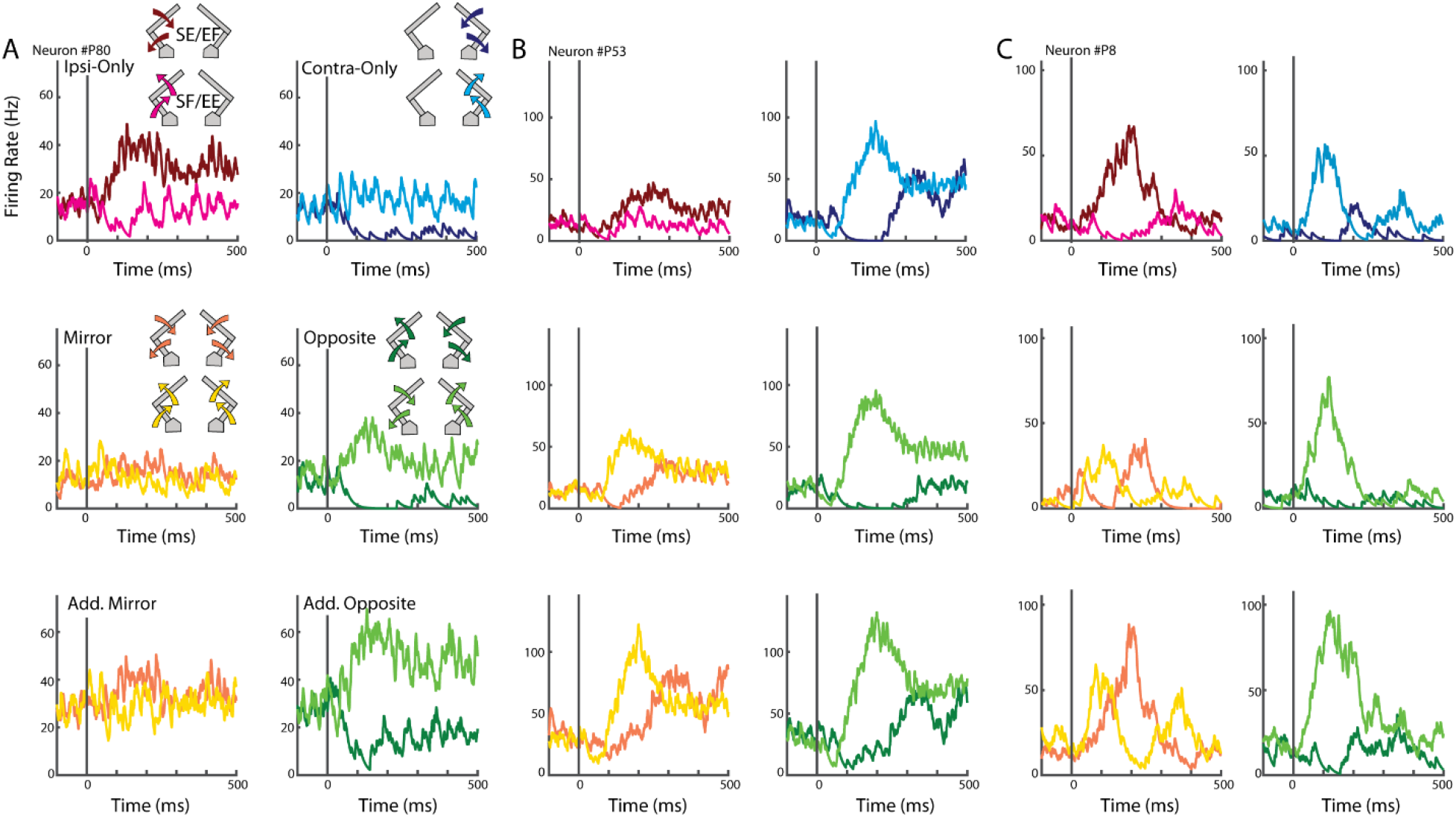
Activities of three example neurons. A) The activities of an example neuron for each load context. Top row: the neuron’s activity for loads applied to the ipsilateral (left ipsi-only) and contralateral (right contra-only) limbs only. For simplicity, only the loads for SE/EF (dark colours) and SF/EE (light colours) are shown. Middle row: the neuron’s activity for the mirror (left) and opposite (right) loads. Bottom row: the expected activities of the neuron if the mirror and opposite activities reflected a linear sum of the contra-only and ipsi-only activities. B-C) Activities from two additional example neurons.

We investigated if a separate population of neurons were active during the unimanual and bimanual contexts. For the ipsi-only and contra-only contexts, we regressed the activity onto the ipsilateral and contralateral loads, respectively. For the bimanual contexts, we concatenated the mirror and opposite contexts and regressed the concatenated activity onto the ipsilateral and contralateral loads. This concatenation was vital as regressing the mirror and opposite contexts separately would result in the ipsilateral loads being collinear to the contralateral loads. Consistent with our previous report (Heming et al., 2019), more neurons had significant fits for the contra-only context than ipsi-only context during the perturbation epoch (Figure 4A,C). We also found a strong overlap between neurons with significant fits for the bimanual and unimanual contexts. For Monkey P/M, 91/74% of neurons had significant fits for the bimanual contexts and at least one of the unimanual contexts (shaded regions). Seven/twelve percent of neurons had significant fits for the unimanual loads only, whereas 2/5% had a significant fit for the bimanual loads only. A similar overlap was observed when we examined the steady-state activity (Figure 4B,D).

**Figure 4:**
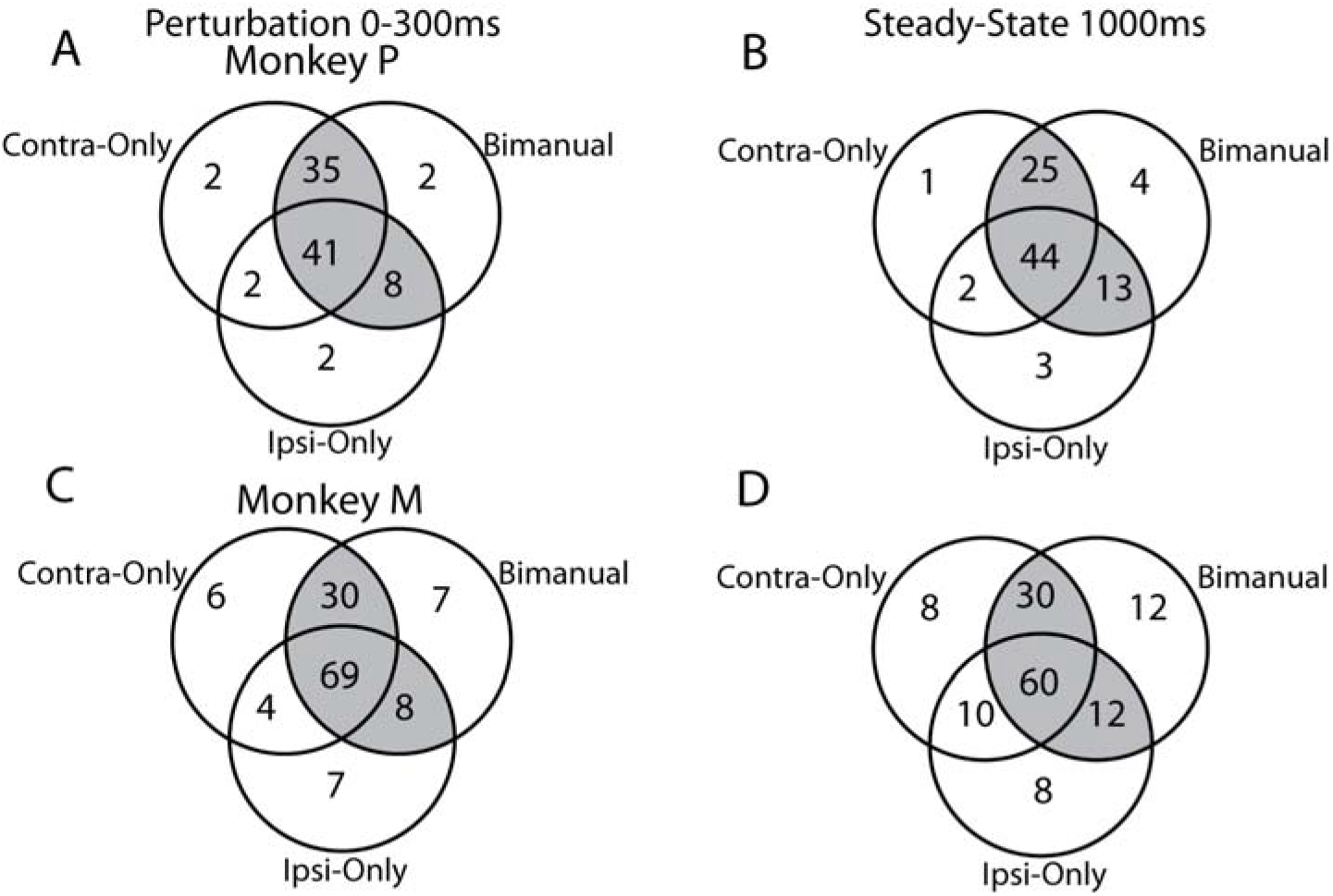
Neuron classification for each load context\. A) For Monkey P, a Venn diagram showing the overlap between neurons with significant fits for the contra-only, ipsi-only and bimanual contexts. Shaded region reflects the neurons with significant fits for at least one of the unimanual (contra-only, ipsi-only) and the bimanual contexts. B) Same as A) for the steady-state epoch. C-D) Same as A-B) for Monkey M.

Next, we investigated if activity during the bimanual context exhibited any suppression relative to the unimanual context. In the perturbation epoch, we estimated each neuron’s dynamic ranges for the mirror and opposite load contexts and compared it with the dynamic ranges from the equivalent additive models. For Monkey P/M, we found the additive mirror model overestimated the activity of 78/83% of neurons (Figure 5A,E), while the additive opposite model overestimated 61/83% of neurons (Figure 5B,F). Across the population, the additive mirror model significantly overestimated the mirror context by 21/49% (Figure 5C,G; paired t-test Monkey P: t(91)=7.4, p<0.001, Monkey M: t(143)=11.3, p<0.001), whereas the additive opposite model overestimated the opposite context by 7/35% (Monkey P: t(91)=2.0, p=0.047, Monkey M: t(143)=10.3 p<0.001). We found a similar overestimation by the additive model when we examined the steady-state epoch (Figure 5D,H; mirror: Monkey P: t(91)=9.0, p<0.001, Monkey M: t(143)=8.4, p<0.001; opposite: Monkey P: t(91)=5.7, p<0.001, Monkey M: t(143)=9.8, p<0.001).

**Figure 5:**
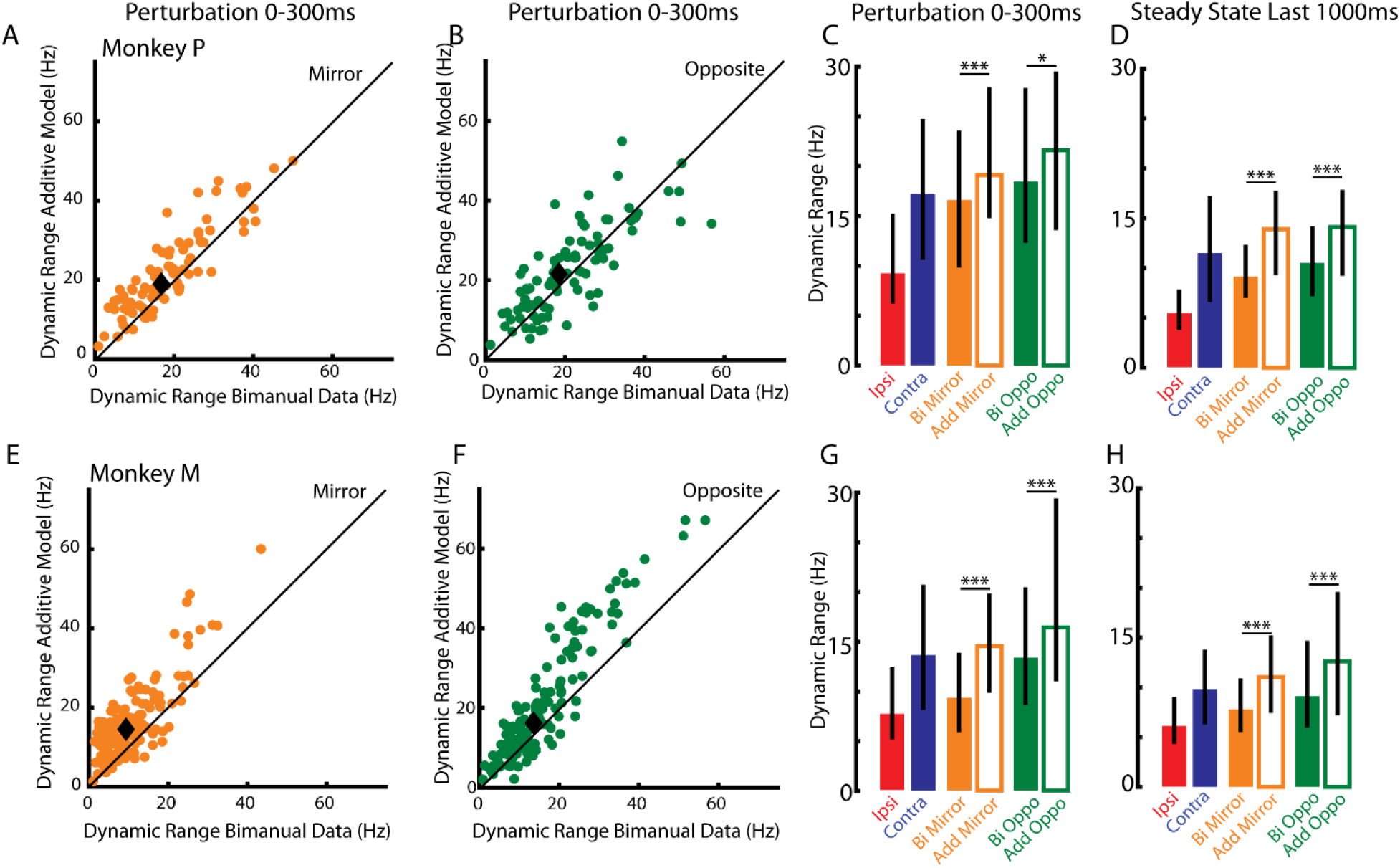
Dynamic range across neurons for the mirror and opposite contexts. A) For Monkey P, comparison between each neuron’s observed dynamic range (abscissa) with its dynamic range predicted by the additive model (ordinate) for the mirror perturbations. Black diamond reflects the median. B) Same as A) for the opposite context. C) The median dynamic range in the perturbation epoch across all recorded neurons (error bars are 25^th^ and 75^th^ percentiles). D) Same as C) for the steady-state epoch. E-H) Same as A-D) for Monkey M. * p<0.05, *** p<0.001.

We explored if the reduction in dynamic range was due to a specific suppression of the ipsilateral-related activity. From the tuning fits, we could separate the activities related to each limb during the bimanual context and calculate the activity magnitudes (see Methods). Figure 6 compares the magnitudes between the unimanual and bimanual contexts for the contralateral- and ipsilateral-related activity. We included only neurons with significant fits for both unimanual and bimanual contexts. In the perturbation epoch, we found the ipsilateral-related activity was smaller during the bimanual context than the unimanual context for 80/65% of neurons in Monkey P/M (Figure 6A, E). Across the population, the ipsilateral-related activity during the bimanual context was 70/82% smaller than the unimanual context for Monkey P/M (Monkey P: paired t-test t(40)=4.9, p<0.001; Monkey M: t(68)=4.1, p<0.001). For the contralateral-related activity of Monkey P, we found the magnitudes of the unimanual and bimanual contexts were similar with almost equal number of neurons residing above and below the unity line (Figure 6B). For Monkey M, the contralateral-related activities were smaller during the bimanual context than the unimanual context for 91% of neurons (Figure 6F). Across the population, the contralateral-related activity during the bimanual context was 79% smaller than the unimanual context (t(68)=8.1, p<0.001). Examining the steady-state activity yielded similar findings (Figure 6C,D,G,H). For both monkeys, we found the activity magnitudes were significantly reduced during the bimanual context for the ipsilateral-related (Monkey P: mean reduction 78%, t(38)=3.8, p<0.001; Monkey M: 83% t(65)=3.2, p=0.002) and contralateral-related activities (Monkey P: 69%, t(38)=5.1, p<0.001; Monkey M: 80%, t(65)=7.0, p<0.001). These data suggest the ipsilateral- and contralateral-related activities exhibited roughly similar levels of suppression.

**Figure 6:**
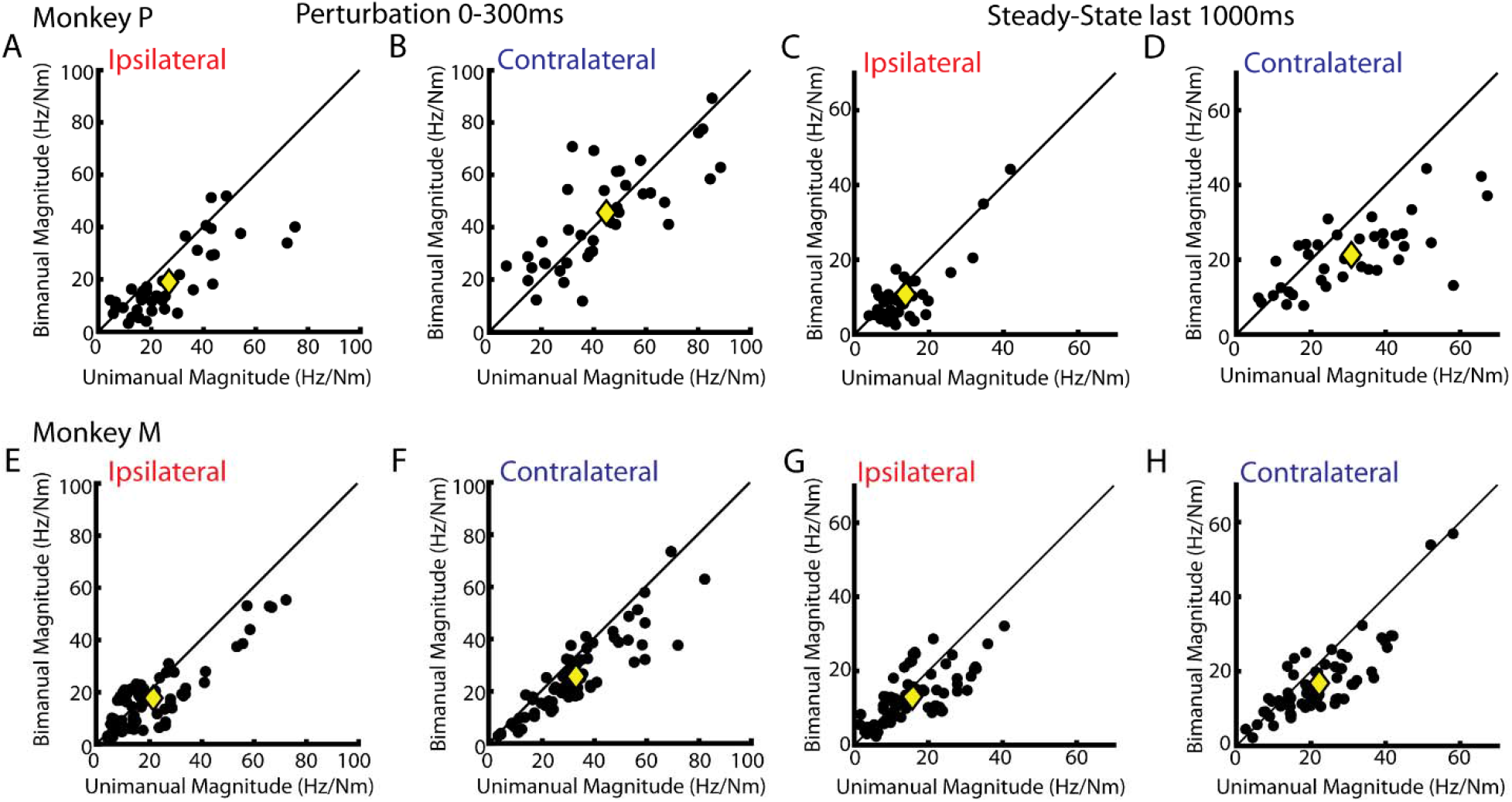
Magnitude changes between the unimanual and bimanual contexts. A) For Monkey P, comparison of the ipsilateral-related magnitudes between the unimanual (abscissa) and bimanual contexts (ordinate) during the perturbation epoch. Yellow diamond indicates the median. B) Same as A) for the contralateral-related magnitudes. C-D) Same as A-B) for the steady-state epoch. E-H) Same as A-D) for Monkey M.

Next, we investigated if the representations changed between unimanual and bimanual contexts. From the tuning fits, we could estimate each neuron’s preferred direction for each limb during the unimanual and bimanual contexts. Figure 7 displays the change in tuning between the unimanual and bimanual contexts. In the perturbation epoch, we found the distribution for the ipsilateral-related activity was centered near the 0° axis indicating that most neurons had similar tuning between the unimanual and bimanual contexts (Figure 7A,E left panel). We quantified how unimodal the distribution was by calculating the Rayleigh (R) coefficient (Figure 7B,F). For comparison, we generated a null distribution where we calculated the change in tuning after shuffling the neurons’ preferred load directions (“shuffle” solid black line, Figure 7B,F). We also generated a distribution that compared the tuning changes expected from two independent samples from the same context (“within context” dashed line). For the ipsilateral-related activity, the change in tuning across contexts was significantly more unimodal (red line, Monkey P/M, Rayleigh coefficient, R = 0.64/0.70) than sampling from a shuffled distribution (both monkeys p<0.001). However, the change in tuning was significantly less unimodal than the within-context distribution (p<0.001), though the difference was small (within context median R = 0.89/0.86). We found similar results when we examined the steady state (Figure 7C,D,G,H).

**Figure 7:**
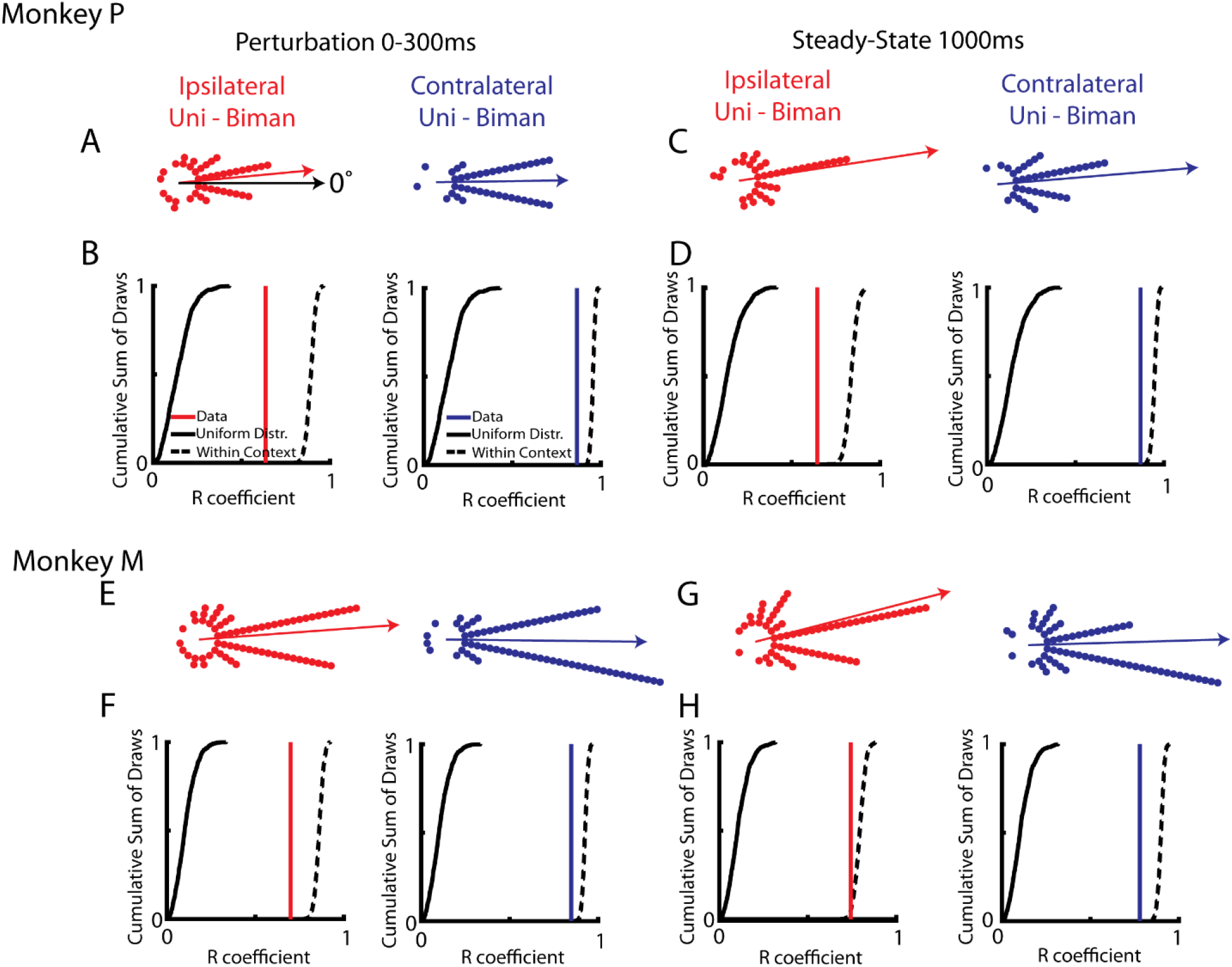
Change of tuning between unimanual and bimanual contexts. A) Polar histograms showing the change in tuning between the unimanual and bimanual contexts for the ipsilateral (left) and contralateral loads (right) during the perturbation epoch. Neurons with no change would lie along the 0° axis. B) Black solid line, cumulative sum of Rayleigh (R) coefficients generated by shuffling neurons and calculating their difference. Black dashed line, cumulative sum of R coefficients generated by comparing the change in tuning within a context. Blue and red lines mark the R coefficients of the data. C-D) Same as A-B) for the steady-state epoch. E-H) Same as A-D) for Monkey M.

For the contralateral-related activity, we also found the distribution was centered near the 0° axis (Figure 7A and E, right panel) and found it was significantly more unimodal (Figure 7B and F right panel blue line, Monkey P/M, R = 0.87/0.85) than sampling from a shuffled distribution (p<0.001). However, the change in tuning was significantly less unimodal than the within-context distribution (p<0.001), though the difference was also small (within context median R = 0.96/0.93). We found similar results when we examined the steady state (Figure 7C,D,G,H).

### Population Analysis

Previously, several groups have shown that ipsilateral- and contralateral-related activities in primary motor cortex could be isolated into orthogonal subspaces during unimanual movements (Ames and Churchland, 2019; Heming et al., 2019). This suggests that motor cortex could in theory maintain the same subspaces for the ipsilateral- and contralateral-related activities during the equivalent bimanual movement. We identified the subspaces for the ipsilateral- and contr4lateral-related activity using the unimanual contexts (Elsayed et al., 2016; Heming et al., 2019). Figure 8A and D, show the variance accounted for (VAF) by the ten dimensions that span the ipsilateral subspace for Monkeys P and M, respectively. For Monkey P/M, this subspace captured 75/60% of the ipsi-only variance and 5/8% of the contra-only variance. This subspace also captured 22/21% of the variance for the mirror context and 31/28% of the variance for the additive mirror model. Similarly, the subspace captured 16/21% of the variance for the opposite context and 34/30% of the variance for the additive opposite model.

**Figure 8:**
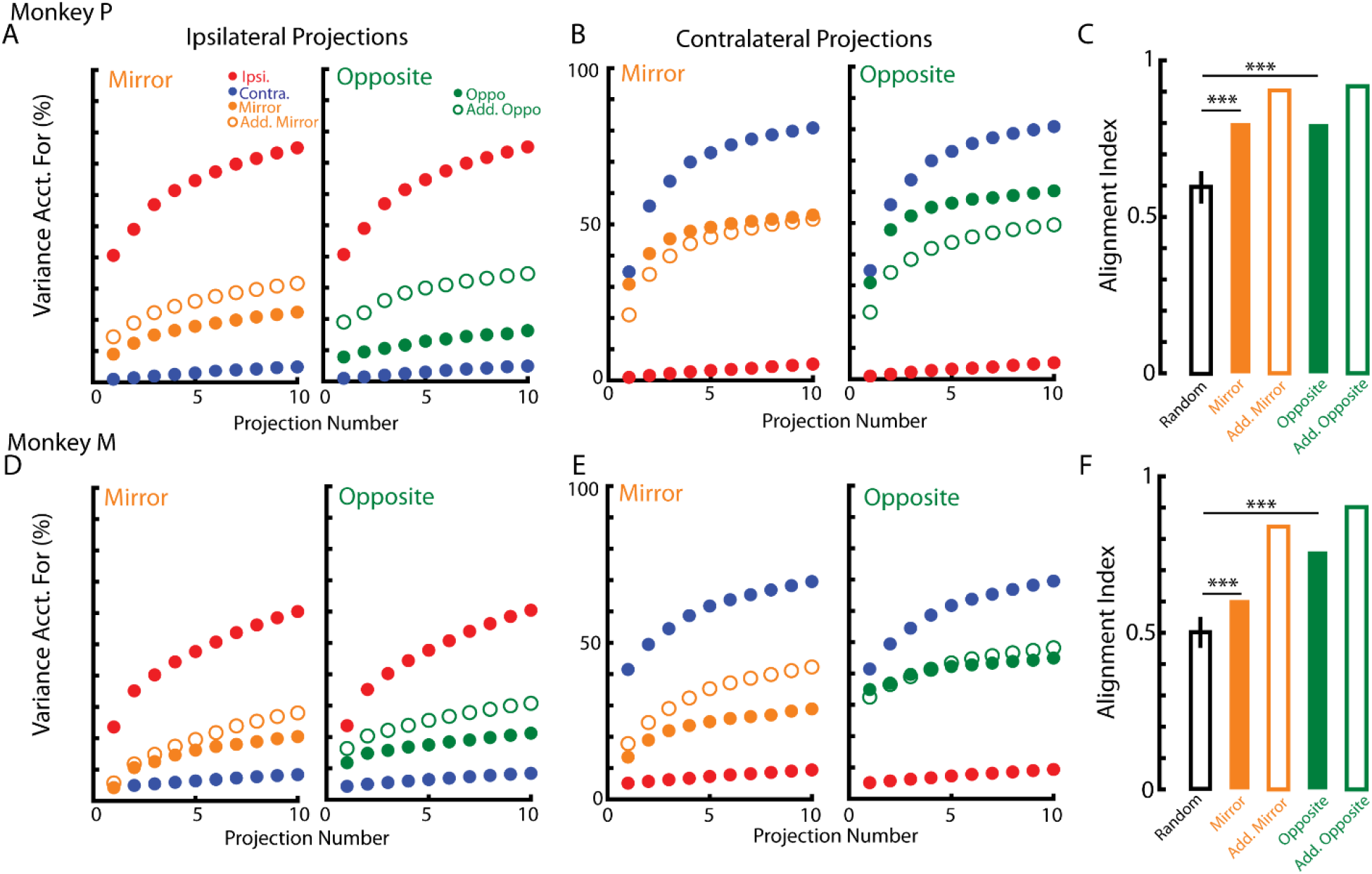
Subspace analysis. A) Left: the variance accounted for by the ipsilateral subspace for the contra-only (blue), ipsi-only (red), mirror (yellow solid), and additive mirror model (yellow open). Right: same as left except the opposite (green solid) and additive opposite model (green open). Note, the contra-only and ipsi-only activities are the same in the left and right panel. Data are plotted as a cumulative sum over the subspace dimensions. B) Same as A) for the contralateral subspace. C) Alignment indices were calculated between the concatenated ipsilateral and contralateral subspaces and the activities for the mirror, opposite and additive models. Random reflects randomly sampling from the data covariance matrix. D-F) Same as A-C) for Monkey M. *** p<0.001.

Figure 8B and E show the VAF by the ten dimensions that span the contralateral subspace for Monkeys P and M, respectively. For Monkey P/M, this subspace captured 80/69% of the contra-only variance (blue dots) and 5/9% of the ipsi-only variance (red dots). This subspace also captured 53/29% of the variance for the mirror context and 52/42% of the variance for the additive mirror model. Similarly, the subspace captured 60/45% of the variance for the opposite context and 49/48% of the variance for the additive opposite model.

We quantified how well the ipsilateral and contralateral subspaces aligned with the subspace that the mirror and opposite activity resided in by calculating the alignment index. The alignment index can range from 0, indicating the subspaces were orthogonal with respect to each other, to 1 indicating complete alignment between the subspaces. A drawback of the alignment index is that including more dimensions in the ipsilateral and contralateral subspaces increases the likelihood that any random subspace will be less orthogonal. We conservatively estimated the alignment index by choosing the top five ipsilateral and contralateral dimensions as most of the neural activity resided in these dimensions. For comparison, we also generated a null distribution that compared how much randomly sampled subspaces were aligned. For both monkeys, the alignment indices for the bimanual contexts (Monkey P/M mirror=0.8/0.6; opposite=0.8/0.76) were lower than the additive model (Monkey P/M mirror=0.9/0.84; opposite=0.92/0.9), however they were significantly greater than the random distribution (p<0.001 for both monkeys, Figure 8C and F). These results suggest that during the bimanual context, a substantial amount of neural activity was maintained in the subspaces identified during the unimanual task.

### Linear vs Nonlinear

Several studies have suggested that representations for the contralateral and ipsilateral limbs are nonlinearly combined during bimanual control (Yokoi et al., 2011; Diedrichsen et al., 2013). We investigated whether nonlinear effects were present in our data by comparing a model with linear terms for the contralateral and ipsilateral loads (linear model) with a model that included linear and nonlinear interaction terms for the contralateral and ipsilateral loads (nonlinear model). Figures 9A and E compares the VAF by the linear (abscissa) and nonlinear models (ordinate) during the perturbation epoch for Monkeys P and M, respectively. We found the linear model captured 89/74% of the variance for Monkey P/M, whereas the nonlinear model captured 93/89% of variance. Also, we found all neurons resided above the unity line consistent with the fact that the nonlinear model had twice as many free parameters. We assessed model performance using Akaikie’s Information Criteria (AIC), which balances how well a model fits with the number of free parameters. Models with lower AIC are preferred to models with larger AIC. Figures 9B and F show the differences between the AIC for the linear and nonlinear models as a cumulative sum for Monkeys P and M, respectively. The cumulative sums reside to the left of the zero line indicating that 97% and 81% of neurons had lower AIC for the linear model than the nonlinear model for Monkeys M and A, respectively. Examining the steady state, we also found the nonlinear model accounted for 6/9% more variance than the linear model for Monkey P/M (Figure 9C,G). However, all neurons had lower AICs for the linear model (Figure 9D,H).

**Figure 9:**
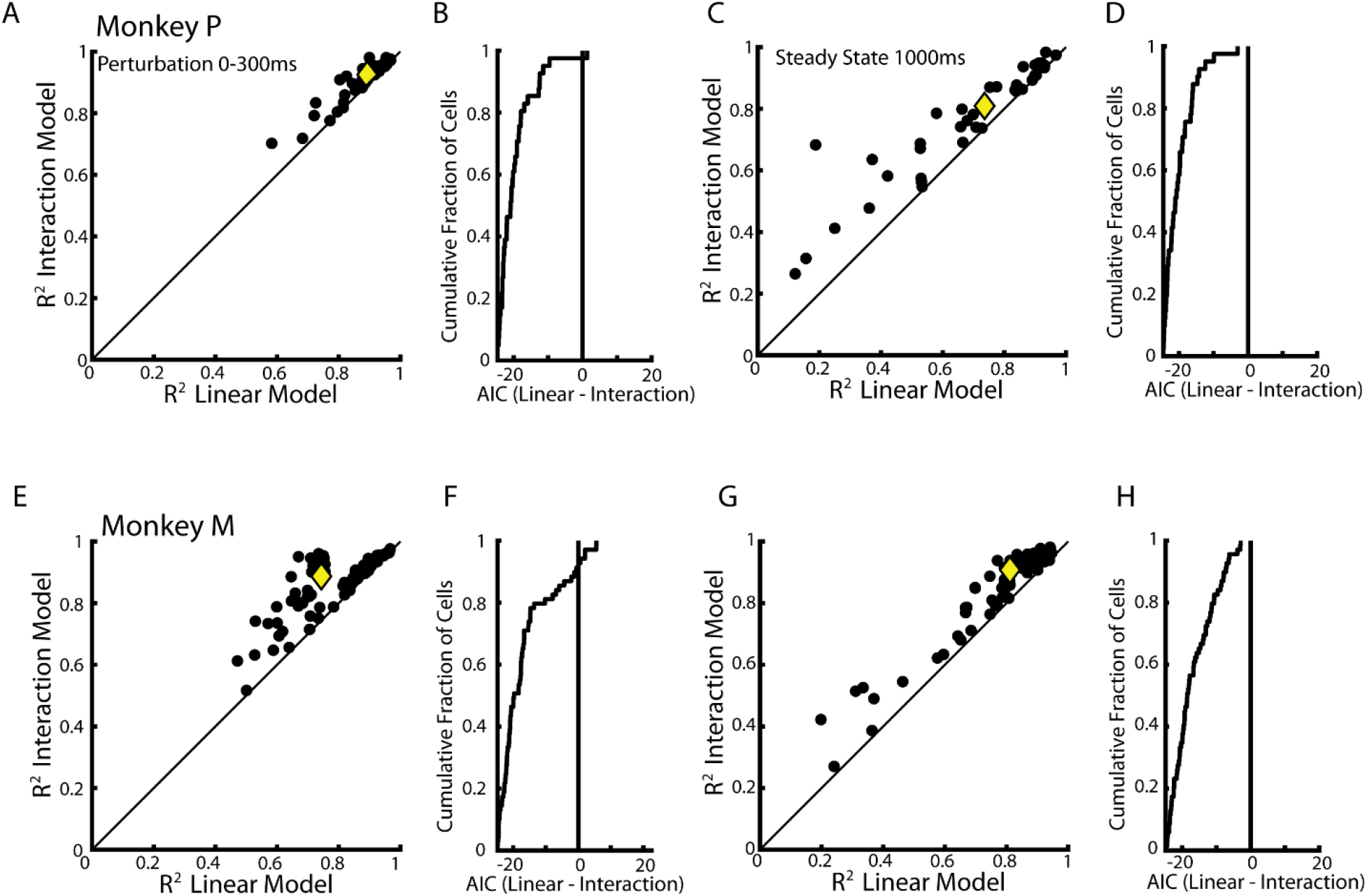
Comparison of the linear and nonlinear models. A) Comparison of model fits between the linear and nonlinear models for each neuron. Yellow diamond reflects the median. B) Difference between the AICs calculated for the linear and nonlinear models. Differences that are less than zero indicate the linear model should be selected, whereas differences greater than zero indicate the nonlinear model should be selected. C-D) Same as A-B) for the steady-state epoch. E-H) Same as A-D) for Monkey M.

## Discussion

We found a substantial overlap between neurons that were responsive to loads applied to either arm (unimanual) and to loads applied to both arms simultaneously (bimanual) in a postural perturbation task. Neurons maintained similar preferred load directions across unimanual and bimanual tasks, but there was a small reduction in activity for the latter. Lastly, we found that the subspace identified for the unimanual loads captured a significant amount of the variance for the bimanual loads. These data highlight how M1 largely maintains its representations of the ipsilateral and contralateral limbs during bimanual control.

Several studies have demonstrated that M1’s representation of the contralateral limb remains stable across time for a given behaviour (Scott and Kalaska, 1997; Chestek et al., 2007; Stevenson et al., 2011). M1 also maintains this representation when adapting to a novel environment (Cherian et al., 2013; Yakovenko and Drew, 2015; Perich and Miller, 2017; Perich et al., 2018; Vyas et al., 2018) and when performing various forms of reaching (Gribble and Scott, 2002; Gallego et al., 2018; Lara et al., 2018). In contrast, large changes in the neural representation have been observed across behavoural tasks (Cheney and Fetz, 1980; Muir and Lemon, 1983; Drew et al., 1996). For example, M1 activity during reaching and locomotion reflect distinct subspaces (Miri et al., 2017). Furthermore, load representations can change dramatically across postural control and reaching, although neurons still maintain similar tuning for external loads across these tasks (Kurtzer et al., 2005; Heming et al., 2016). Thus, neural representations in M1 remain relatively constant for a given behaviour but can show substantial changes across behaviours.

Here, we found the contralateral representation remains stable across unimanual and bimanual contexts. We found a reduction of activity that may reflect a corresponding reduction in the motor output. We cannot rule this out as we did not record muscle activity, but hand kinematics were similar between unimanual and bimanual loads for the first 300ms after the load was applied. Furthermore, we observed a similar reduction during the steady-state epoch when motor output should be comparable between the unimanual and bimanual loads. Importantly, the preferred load directions remained quite constant, and the subspace identified during the unimanual task captured as much of the variance during the bimanual task as expected from the additive model. Thus, there was a small reduction in activity, but the basic pattern of activity across behavioural contexts remained stable. Similar results were generally found for the ipsilateral representation, although the ipsilateral subspace captured only 66% of the activity during the bimanual loads as compared to the additive model. Thus, while a substantive proportion of the representation was maintained it was less than that observed for the contralateral limb. The ability to simultaneously represent both limbs while performing bimanual motor actions may reflect that the subspaces associated with each limb were orthogonal.

In contrast, premotor cortical regions show a greater change in neural representations between unimanual and bimanual motor actions (Tanji et al., 1987, 1988; Rokni et al., 2003; Willett et al., 2020). During bimanual movements, Willett and colleagues, (2019) found relatively small reductions in the contralateral representation in the premotor cortex of humans, but larger reductions for the ipsilateral representation on the order of 50%. Interestingly, they found that the ipsilateral and contralateral representations were in subspaces that overlapped more than M1’s representations. Cisek et al., (2003) also found that the preferred direction of neurons during reaching are correlated for the two limbs. This lack of orthogonality in premotor regions may result in a reduction of the ipsilateral representation in order to reduce interference during bimanual motor actions (Rokni et al., 2003; Willett et al., 2020).

Although speculative, these differences in the organization of ipsilateral and contralateral representations may reflect the types of information that are represented in these cortical areas. Studies have highlighted that premotor cortical activity is more related to extrinsic features of motor actions, whereas M1 activity is more related to intrinsic features related to the motor periphery (Evarts, 1968; Humphrey, 1972; Cheney and Fetz, 1980; Fromm, 1983; Werner et al., 1991; Scott and Kalaska, 1997; Scott et al., 1997; Shen and Alexander, 1997a, 1997b). It may be that goal-related features of a task are more broadly reflected across the entire premotor network. In natural situations, this broad expression of the behavioural goal may prove valuable in order to permit rapid alternate motor strategies to attain the goal, such as using the other limb to reach and grasp an object of interest. In contrast, when there are independent goals for different motor effectors the premotor representation of the goal associated with the appropriate effector is maintained while the other goal representations are suppressed. In contrast, M1 activity is more related to the details of motor execution which is more effector specific and M1 is also closer to downstream motor targets. Thus, M1 exhibits independent representations of the two limbs, but this allows both representations to be maintained during bimanual motor actions.

Previous studies by Vaadia and colleagues had explored bimanual coordination in M1 (Steinberg et al., 2002; Rokni et al., 2003). However, their population of neurons exhibited functional properties more similar to premotor cortex. They found neurons had similar tuning for the contralateral and ipsilateral limbs during unimanual reaches (Steinberg et al., 2002). They also found a substantial change in a neuron’s preferred direction and an ~50% reduction in magnitude for the ipsilateral-related activity between unimanual and bimanual reaches (Rokni et al., 2003). This may reflect some fluidity in ipsilateral representations across animals or behavioural tasks, postural versus reaching. It is also possible that their M1 recordings were from the transition zone between premotor cortex and M1 which exhibits properties reflecting a mixture of the two areas (Cisek et al., 2003).

We used floating micro-electrode arrays to record from M1 that was positioned on the surface of the precentral gyrus. As a result, we did not sample from the most caudal portion of M1 which lies in the bank of the central sulcus. Studies have suggested a rostral-caudal gradient across motor cortex for several attributes. The caudal motor cortex exhibits greater number of cortico-motor neurons (Rathelot and Strick, 2009; Witham et al., 2016), greater independence of tuning between the upper limbs (Cisek et al., 2003), decreased preparatory activity (Crammond and Kalaska, 2000) and greater steady-state activity during postural control (Crammond and Kalaska, 1996) than rostral motor cortex (i.e. premotor cortex). If a gradient does exist, then caudal M1 likely also maintains orthogonal subspaces for the ipsilateral and contralateral limbs but may show even less reduction in activity during bimanual motor tasks than rostral M1.

The parietal reach region (PRR) also displays neural representations related to motor actions of both limbs (Kermadi et al., 2000; Chang et al., 2008; Mooshagian et al., 2018). PRR is primarily involved with controlling the contralateral limb (Chang et al., 2008; Yttri et al., 2013), however neurons in PRR respond prior to movements of the contralateral and ipsilateral limbs as well as upcoming saccades (Chang et al., 2008; Chang and Snyder, 2012). However, this ipsilateral activity is predominantly related to a sensory response to the visual target, whereas responses for the contralateral limb are related to both the sensory event and motor planning (Mooshagian et al., 2018).

It is not clear whether representing both limbs by one hemisphere and the change to these representations during bimanual motor actions influences actual motor function. Given the behavioural goal was identical for a given limb during unimanual and bimanual tasks, one might expect that any change in the neural representations might impact control. As stated above, we did not observe substantive changes in the kinematics of movement in this relatively simple postural perturbation task. However, the motor system appears to prefer mirror symmetric movements of the limb even when instructed to perform anti-symmetric movements (Kelso, 1984). Furthermore, learning a force field while performing a unimanual reach only partially transfers to the equivalent bimanual reach (Nozaki et al., 2006; Nozaki and Scott, 2009; Howard et al., 2010). These observations may reflect interactions between the ipsilateral and contralateral representations in motor cortex during bimanual motor tasks.

The presence of bimanual representations in motor cortex may support bimanual coordination in tasks when the two limbs work together to perform a common goal. Currently, most neurophysiological investigations of bimanual control, including our own, have utilized tasks where the goals of each limb are independent, thus requiring minimal interlimb coordination (Donchin et al., 1998; Steinberg et al., 2002; Rokni et al., 2003; Willett et al., 2020). Future studies should investigate behaviours that require interlimb coordination to attain a common goal (Diedrichsen et al., 2004; Dimitriou et al., 2012; Córdova Bulens et al., 2017). In these contexts, sensory feedback from one limb can elicit goal-directed motor actions in the opposite limb in ~70ms (Diedrichsen, 2007; Mutha and Sainburg, 2009; Omrani et al., 2013). It is likely that these interlimb feedback responses involve interactions between overlapping subspaces in motor cortex.

## Acknowledgements

We thank Kim Moore, Simone Appaqaq, Justin Peterson, and Helen Bretzke for their laboratory and technical assistance and members of the LIMB lab for constructive criticisms. This work was supported by grants from the Canadian Institute of Health Research. KPC was supported by an OGS scholarship. EAH was supported by an NSERC scholarship. SHS was supported by a GSK chair in Neuroscience.

## References

Ames KC, Churchland MM (2019) Motor cortex signals for each arm are mixed across hemispheres and neurons yet partitioned within the population response. eLife 8:e46159.

Batschelet E (1981) Circular Statistics in Biology. New York: Academic Press.

Berlot E, Prichard G, O’Reilly J, Ejaz N, Diedrichsen J (2019) Ipsilateral finger representations in the sensorimotor cortex are driven by active movement processes, not passive sensory input. J Neurophysiol 121:418–426.

Brosamle C, Schwab ME (1997) Cells of origin, course, and termination patterns of the ventral, uncrossed component of the mature rat corticospinal tract. J Comp Neurol 386:293–303.

Burnham KP, Anderson DR (2004) Model selection and multimodel inference: A practical infromation-theoretic approach, 2nd ed. New York, NY: Springer-Verlag.

Chang SWC, Dickinson AR, Snyder LH (2008) Limb-Specific Representation for Reaching in the Posterior Parietal Cortex. J Neurosci 28:6128–6140.

Chang SWC, Snyder LH (2012) The representations of reach endpoints in posterior parietal cortex depend on which hand does the reaching. J Neurophysiol 107:2352–2365.

Cheney PD, Fetz EE (1980) Functional classes of primate corticomotoneuronal cells and their relation to active force. J Neurophysiol 44:773–791.

Cherian A, Fernandes HL, Miller LE (2013) Primary motor cortical discharge during force field adaptation reflects muscle-like dynamics. J Neurophysiol 110:768–783.

Chestek CA, Batista AP, Santhanam G, Yu BM, Afshar A, Cunningham JP, Gilja V, Ryu SI, Churchland MM, Shenoy KV (2007) Single-Neuron Stability during Repeated Reaching in Macaque Premotor Cortex. J Neurosci 27:10742–10750.

Cisek P, Crammond DJ, Kalaska JF (2003) Neural Activity in Primary Motor and Dorsal Premotor Cortex In Reaching Tasks With the Contralateral Versus Ipsilateral Arm. J Neurophysiol 89:922–942.

CÓrdova Bulens D, Crevecoeur F, Thonnard J-L, Lefèvre P (2017) Optimal use of limb mechanics distributes control during bimanual tasks. J Neurophysiol:jn.00371.2017.

Cramer SC, Finklestein SP, Schaechter JD, Bush G, Rosen BR (1999) Activation of Distinct Motor Cortex Regions During Ipsilateral and Contralateral Finger Movements. J Neurophysiol 81:383–387.

Crammond DJ, Kalaska JF (1996) Differential relation of discharge in primary motor cortex and premotor cortex to movements versus actively maintained postures during a reaching task. Exp Brain Res 108 Available at: http://link.springer.com/10.1007/BF00242903 [Accessed March 1, 2019].

Crammond DJ, Kalaska JF (2000) Prior Information in Motor and Premotor Cortex: Activity During the Delay Period and Effect on Pre-Movement Activity. J Neurophysiol 84:986–1005.

Diedrichsen J (2007) Optimal Task-Dependent Changes of Bimanual Feedback Control and Adaptation. Curr Biol 17:1675–1679.

Diedrichsen J, Nambisan R, Kennerley SW, Ivry RB (2004) Independent on-line control of the two hands during bimanual reaching. Eur J Neurosci 19:1643–1652.

Diedrichsen J, Wiestler T, Krakauer JW (2013) Two Distinct Ipsilateral Cortical Representations for Individuated Finger Movements. Cereb Cortex 23:1362–1377.

Dimitriou M, Franklin DW, Wolpert DM (2012) Task-dependent coordination of rapid bimanual motor responses. J Neurophysiol 107:890–901.

Donchin O, Gribova A, Steinberg O, Bergman H, Vaadia E (1998) Primary motor cortex is involved in bimanual coordination. Nature 395:274–278.

Downey JE, Quick KM, Schwed N, Weiss JM, Wittenberg GF, Boninger ML, Collinger JL (2019) Primary motor cortex has independent representations for ipsilateral and contralateral arm movements but correlated representations for grasping. medRxiv Available at: http://medrxiv.org/lookup/doi/10.1101/19008128 [Accessed October 28, 2019].

Drew T, Jiang W, Kably B, Lavoie S (1996) Role of the motor cortex in the control of visually triggered gait modifications. Can J Physiol Pharmacol 74:426–442.

Dum RP, Strick PL (1996) Spinal Cord Terminations of the Medial Wall Motor Areas in Macaque Monkeys. J Neurosci 16:6513–6525.

Elsayed GF, Lara AH, Kaufman MT, Churchland MM, Cunningham JP (2016) Reorganization between preparatory and movement population responses in motor cortex. Nat Commun 7 Available at: http://www.nature.com/articles/ncomms13239 [Accessed November 26, 2018].

Evarts EV (1968) Relation of pyramidal tract activity to force exterted durin voluntary movement. J Neurophysiol 31:14–27.

Fraser GW, Schwartz AB (2012) Recording from the same neurons chronically in motor cortex. J Neurophysiol 107:1970–1978.

Fromm C (1983) Changes of steady state activity in motor cortex consistent with the length-tension relation of muscle. Pflugers Arch 398:318–323.

Gallego JA, Perich MG, Naufel SN, Ethier C, Solla SA, Miller LE (2018) Cortical population activity within a preserved neural manifold underlies multiple motor behaviors. Nat Commun 9 Available at: http://www.nature.com/articles/s41467-018-06560-z [Accessed November 16, 2018].

Ganguly K, Secundo L, Ranade G, Orsborn A, Chang EF, Dimitrov DF, Wallis JD, Barbaro NM, Knight RT, Carmena JM (2009) Cortical Representation of Ipsilateral Arm Movements in Monkey and Man. J Neurosci 29:12948–12956.

Gribble PL, Scott SH (2002) Overlap of internal models in motor cortex for mechanical loads during reaching. Nature 417:938–941.

Heming EA, Cross KP, Takei T, Cook DJ, Scott SH (2019) Independent representations of ipsilateral and contralateral limbs in primary motor cortex. eLife 8:e48190.

Heming EA, Lillicrap TP, Omrani M, Herter TM, Pruszynski JA, Scott SH (2016) Primary motor cortex neurons classified in a postural task predict muscle activation patterns in a reaching task. J Neurophysiol 115:2021–2032.

Herter TM, Korbel T, Scott SH (2009) Comparison of Neural Responses in Primary Motor Cortex to Transient and Continuous Loads During Posture. J Neurophysiol 101:150–163.

Howard IS, Ingram JN, Wolpert DM (2010) Context-Dependent Partitioning of Motor Learning in Bimanual Movements. J Neurophysiol 104:2082–2091.

Humphrey DR (1972) Relating motor cortex spike trains to measures of motor performance. Brain Res 40:7–18.

Kelso JAS (1984) Phase transitions and critical behaviour in human bimanual coordination. Am J Physiol 246: R1000–R1004.

Kermadi I, Calciati T, Rouiller EM (1998) Neuronal activity in the primate supplementary motor area and the primary motor cortex in relation to spatio-temporal bimanual coordination. Somatosens Mot Res 15:287–308.

Kermadi I, Liu Y, Rouiller EM (2000) Do bimanual motor actions involve the dorsal premotor (PMd), cingulate (CMA) and posterior parietal (PPC) cortices? Comparison with primary and supplementary motor cortical areas. Somatosens Mot Res 17:255–271.

Kurtzer I, Herter TM, Scott SH (2005) Random change in cortical load representation suggests distinct control of posture and movement. Nat Neurosci 8:498–504.

Kuypers HGJM (2011) Anatomy of the Descending Pathways. In: Comprehensive Physiology, pp 597–666. American Cancer Society. Available at: https://onlinelibrary.wiley.com/doi/abs/10.1002/cphy.cp010213 [Accessed November 20, 2019]

Lacroix S, Havton LA, McKay H, Yang H, Brant A, Roberts J, Tuszynski MH (2004) Bilateral corticospinal projections arise from each motor cortex in the macaque monkey: A quantitative study. J Comp Neurol 473:147–161.

Lara AH, Elsayed GF, Zimnik AJ, Cunningham JP, Churchland MM (2018) Conservation of preparatory neural events in monkey motor cortex regardless of how movement is initiated. eLife 7:e31826.

Miri A, Warriner CL, Seely JS, Elsayed GF, Cunningham JP, Churchland MM, Jessell TM (2017) Behaviorally Selective Engagement of Short-Latency Effector Pathways by Motor Cortex. Neuron 95:683-696.e11.

Montgomery LR, Herbert WJ, Buford JA (2013) Recruitment of ipsilateral and contralateral upper limb muscles following stimulation of the cortical motor areas in the monkey. Exp Brain Res 230:153–164.

Mooshagian E, Wang C, Holmes CD, Snyder LH (2018) Single Units in the Posterior Parietal Cortex Encode Patterns of Bimanual Coordination. Cereb Cortex N Y NY 28:1549–1567.

Muir RB, Lemon RN (1983) Corticospinal neurons with a special role in precision grip. Brain Res 261:312–316.

Mutha PK, Sainburg RL (2009) Shared Bimanual Tasks Elicit Bimanual Reflexes During Movement. J Neurophysiol 102:3142–3155.

Nozaki D, Kurtzer I, Scott SH (2006) Limited transfer of learning between unimanual and bimanual skills within the same limb. Nat Neurosci 9:1364–1366.

Nozaki D, Scott SH (2009) Multi-compartment model can explain partial transfer of learning within the same limb between unimanual and bimanual reaching. Exp Brain Res 194:451–463.

Omrani M, Diedrichsen J, Scott SH (2013) Rapid feedback corrections during a bimanual postural task. J Neurophysiol 109:147–161.

Perich MG, Gallego JA, Miller L (2018) A Neural Population Mechanism For Rapid Learning. Neuron 100:964–976.

Perich MG, Miller LE (2017) Altered tuning in primary motor cortex does not account for behavioral adaptation during force field learning. Exp Brain Res 235:2689–2704.

Pruszynski JA, Omrani M, Scott SH (2014) Goal-Dependent Modulation of Fast Feedback Responses in Primary Motor Cortex. J Neurosci 34:4608–4617.

Rathelot J-A, Strick PL (2009) Subdivisions of primary motor cortex based on cortico-motoneuronal cells. Proc Natl Acad Sci 106:918–923.

Rokni U, Steinberg O, Vaadia E, Sompolinsky H (2003) Cortical Representation of Bimanual Movements. J Neurosci 23:11577–11586.

Rosenzweig ES, Brock JH, Culbertson MD, Lu P, Moseanko R, Edgerton VR, Havton LA, Tuszynski MH (2009) Extensive spinal decussation and bilateral termination of cervical corticospinal projections in rhesus monkeys. J Comp Neurol 513:151–163.

Scott SH (1999) Apparatus for measuring and perturbing shoulder and elbow joint positions and torques during reaching. J Neurosci Methods 89:119–127.

Scott SH, Kalaska JF (1997) Reaching Movements With Similar Hand Paths But Different Arm Orientations. I. Activity of Individual Cells in Motor Cortex. J Neurophysiol 77:826–852.

Scott SH, Sergio LE, Kalaska JF (1997) Reaching Movements With Similar Hand Paths but Different Arm Orientations. II. Activity of Individual Cells in Dorsal Premotor Cortex and Parietal Area 5. J Neurophysiol 78:2413–2426.

Shen L, Alexander GE (1997a) Preferential Representation of Instructed Target Location Versus Limb Trajectory in Dorsal Premotor Area. J Neurophysiol 77:1195–1212.

Shen L, Alexander GE (1997b) Neural Correlates of a Spatial Sensory-To-Motor Transformation in Primary Motor Cortex. J Neurophysiol 77:1171–1194.

Soteropoulos DS, Edgley SA, Baker SN (2011) Lack of Evidence for Direct Corticospinal Contributions to Control of the Ipsilateral Forelimb in Monkey. J Neurosci 31:11208–11219.

Steinberg O, Donchin O, Gribova A, De Oliveira SC, Bergman H, Vaadia E (2002) Neuronal populations in primary motor cortex encode bimanual arm movements: Population vectors in bimanual movements. Eur J Neurosci 15:1371–1380.

Stevenson IH, Cherian A, London BM, Sachs NA, Lindberg E, Reimer J, Slutzky MW, Hatsopoulos NG, Miller LE, Kording KP (2011) Statistical assessment of the stability of neural movement representations. J Neurophysiol 106:764–774.

Tanji J, Okano K, Sato KC (1987) Relation of neurons in the nonprimary motor cortex to bilateral hand movement. Nature 327:618–620.

Tanji J, Okano K, Sato KC (1988) Neuronal activity in cortical motor areas related to ipsilateral, contralateral, and bilateral digit movements of the monkey. J Neurophysiol 60:325–343.

Thompson KG, Hanes DP, Bichot NP, Schall JD (1996) Perceptual and motor processing stages identified in the activity of macaque frontal eye field neurons during visual search. J Neurophysiol 76:4040–4055.

Vyas S, Even-Chen N, Stavisky SD, Ryu SI, Nuyujukian P, Shenoy KV (2018) Neural Population Dynamics Underlying Motor Learning Transfer. Neuron 97:1177–1186.e3.

Werner W, Bauswein E, Fromm C (1991) Static firing rates of premotor and primary motor cortical neurons associated with torque and joint position. Exp Brain Res 86:293–302.

Willett FR, Deo DR, Avansino DT, Rezaii P, Hochberg LR, Henderson JM, Shenoy KV (2020) Hand Knob Area of Premotor Cortex Represents the Whole Body in a Compositional Way. Cell 0 Available at: https://www.cell.com/cell/abstract/S0092-8674(20)30220-8 [Accessed March 26, 2020].

Witham CL, Fisher KM, Edgley SA, Baker SN (2016) Corticospinal Inputs to Primate Motoneurons Innervating the Forelimb from Two Divisions of Primary Motor Cortex and Area 3a. J Neurosci 36:2605–2616.

Yakovenko S, Drew T (2015) Similar Motor Cortical Control Mechanisms for Precise Limb Control during Reaching and Locomotion. J Neurosci 35:14476–14490.

Yokoi A, Hirashima M, Nozaki D (2011) Gain Field Encoding of the Kinematics of Both Arms in the Internal Model Enables Flexible Bimanual Action. J Neurosci 31:17058–17068.

Yttri EA, Wang C, Liu Y, Snyder LH (2013) The parietal reach region is limb specific and not involved in eye-hand coordination. J Neurophysiol 111:520–532.

